# Integrin promotes basement membrane crossing and branching of an invading intracellular tube

**DOI:** 10.1101/2025.04.24.650450

**Authors:** Lauren N. Meyer, Michael Hertel, Jeremy Nance

**Author notes:** Author for correspondence: Jeremy Nance, University of Wisconsin – Madison, 1525 Linden Drive, 321 Bock Labs, Madison, WI 53706.

## Abstract

The narrowest biological tubes are comprised of cells that hollow to form an intracellular lumen. Here, we examine early lumenogenesis of the *C. elegans* excretory cell, which branches to form an H-shaped intracellular tube spanning the length of the worm. Using genetically paralyzed embryos to freeze movement, we describe lumen initiation and branching for the first time using time-lapse fluorescence microscopy. We show that the excretory cell lumen forms through a plasma membrane invasion mechanism when a nascent lumen grows from the plasma membrane into the cytoplasm. The lumen subsequently extends along the left-right axis before branching to form anterior-posterior projections. Through a genetic screen, we identify mutations in *ina-1/⍺-integrin* and *pat-3/β-integrin* that block lumenogenesis at the anterior-posterior branching step, and we show that integrin function is required within the excretory cell. Finally, we find that the excretory cell crosses the epidermal basement membrane where anterior-posterior branches form and demonstrate that basement membrane crossing fails in integrin mutant embryos. Our findings reveal how an intracellular lumen initiates and branches and identify integrins and basement membrane as key branching regulators.

## INTRODUCTION

Most organs contain networks of multicellular tubes that vary widely in shape and diameter. For example, tubes present in the human respiratory tract range from large bronchi, which carry air into and out of the lungs, to small sac-like alveoli, which exchange gases brought into the lung with the vasculature (Banavar et al., 2024). The cells lining multicellular tubes are epithelial in nature; their apical surfaces face the tube lumen and basolateral surfaces interact with other cells and the extracellular matrix (Camelo and Luschnig, 2021; Shaye and Soto, 2021). Cells lining the tube lumen attach to each other with adherens junctions and tight junctions, which seal the lumen and provide mechanical connections important for tube morphogenesis and integrity.

The smallest-bore tubes, which run through the cytoplasm of individual cells and are collectively called intracellular tubes, form using fundamentally different mechanisms (Sundaram and Cohen, 2017). Some intracellular tubes arise when a cell wraps around an extracellular space and attaches back to itself, thereby enclosing the circumscribed extracellular space as a lumen (Rasmussen et al., 2008; Stone et al., 2009). Others develop via a plasma membrane invasion mechanism, when one or more apical membrane domains invades the cytoplasm to hollow the cell center (Dong et al., 2009; Gervais and Casanova, 2010). Intracellular tubes can connect to other tubes at both ends, as occurs in the ascidian notochord and smallest vertebrate blood vessels (Dong et al., 2009; Herwig et al., 2011; Lenard et al., 2013). Alternatively, they can be blind-ended, as in *Drosophila* terminal tracheal cells (Gervais and Casanova, 2010).

The *C. elegans* excretory system, which functions in osmoregulation (Nelson and Riddle, 1984), provides a genetic model system for the study of intracellular tubulogenesis (Sundaram and Buechner, 2016). The excretory system is composed of connected intracellular tubes from three cells – the pore cell, the duct cell, and the excretory cell – as well as an interacting gland cell (Nelson et al., 1983). The lumen of the pore cell connects to surface epidermal cells, creating an opening for the excretory system on the worm’s ventral surface. The pore cell lumen connects to the duct cell lumen, which in turn connects to the excretory cell lumen. The excretory cell branches to form four lumenized blind-ended projections called ‘canals’, which run along the left and right sides of the worm. The gland cell, whose function is unknown, interfaces with the excretory system lumen at the junction of the excretory and duct cells, forming a tricellular tight junction at this site (Nelson et al., 1983).

Remarkably, the three intracellular tubes of the excretory system form using different methods of tubulogenesis. The pore cell and duct cell lumenize via the wrapping mechanism, creating an auto-cellular junction when they reattach to themselves (Stone et al., 2009); the pore cell auto-cellular junction remains, creating a ‘seamed’ intracellular tube, whereas the duct cell auto-cellular junction disappears as a result of AFF-1-mediated self-fusion (Soulavie et al., 2018). By contrast, the excretory cell forms an intracellular lumen without wrapping. The excretory cell body takes on its complex shape during embryogenesis (Buechner et al., 1999; Hedgecock et al., 1990; Hedgecock et al., 1987). The cell first extends bilaterally towards the left and right sides of the embryo. Subsequently, its left and right extensions split to form anterior and posterior projections that grow between the epidermis and the epidermal basement membrane; extension of the excretory projections continues through the first larval stage, at which point the posterior projections reach nearly to the end of the body. As the excretory cell projections grow, the cell hollows to form a lumen that mirrors the pattern of the projections, creating the mature excretory canals. Large canalicular vesicles fuse and make stable connections with the canal lumen, creating a Swiss-cheese pattern that increases lumen surface area (Nelson et al., 1983).

To date, studies examining how the excretory cell lumenizes have focused largely on cytoskeletal regulators and polarized trafficking pathways, which are needed for lumen growth and maintenance (Shaye and Soto, 2021; Sundaram and Buechner, 2016; Sundaram and Cohen, 2017). However, the initial stages of lumen formation, including lumen initiation and branching – remain relatively unexplored. In contrast to the branches of *Drosophila* terminal tracheal cells, which do not form reproducible patterns even among homologous cells on the left and right sides of the body (Gavrilchenko et al., 2024), the excretory cell lumen branches form at the same position within the developing embryo. Because the early stages of lumen growth and branching occur after the embryo begins to move vigorously within the eggshell, which complicates live image analysis, these events have been examined only in fixed embryos using electron microscopy. Two important questions remain unresolved that could be more conclusively addressed using timelapse microscopy. First, it is unclear how the lumen initiates and connects to the duct cell lumen; different analyses of embryo electron micrographs have suggested alternative mechanisms – hollowing of the cytoplasm by vacuole formation (Berry et al., 2003) or invasion of the apical membrane into the cytoplasm (Stone et al., 2009). Second, it is not known how the excretory cell lumen branches in a reproducible pattern. Since anterior-posterior branching occurs at the lateral sides of the embryo adjacent to the epidermis, interactions between the excretory cell and epidermal cells or the epidermal basement membrane may contribute to this event. These hypotheses have not been explored.

Here, we investigate the mechanisms of excretory cell lumen formation and branching using live imaging and mutant analysis. We first detect the lumen at the interface between the excretory cell and the adjacent duct, suggesting that the lumen initiates at this site and grows into the cell. By genetically paralyzing embryos to halt their movement within the eggshell, we visualize lumen branching in living embryos for the first time and demonstrate a role for integrin signaling in anterior-posterior branch formation. Finally, we show that the excretory cell extensions cross the epidermal basement membrane before branching and fail to do so in the absence of integrin function, suggesting that integrin-mediated crossing of the basement membrane is required for branching and migration of the excretory cell canals.

## RESULTS

### The excretory cell lumen invades inward from the site of duct cell contact

The excretory cell lumen appears during mid-embryogenesis after the excretory, duct, and pore cells converge (Stone et al., 2009). To visualize the initial stages of lumen growth in living embryos, we created transgenic lines that express membrane-targeted YFP (YFP fused to the PIP_2_-binding PH domain of PLC∂1; hereafter Exc^YFP-Mem^) from a *lin-3* promotor fragment known to drive expression in the excretory cell (Abdus-Saboor et al., 2011; Hwang and Sternberg, 2004). Exc^YFP-Mem^ was expressed in the excretory cell beginning soon after its birth and persisted until early larval stages. Initially, Exc^YFP-Mem^ marked the plasma membrane of the excretory cell uniformly (Fig. 1B; n = 21 embryos); one or more transient, filopodia-like extensions of plasma membrane were often present on the anterior side of the cell at this stage (e.g. Fig. 1B,C,E’). Beginning at the 1.5-fold stage, Exc^YFP-Mem^ developed a circular enrichment on the ventral side of the excretory cell (Fig. 1C, arrowhead; n = 45 embryos). By the 2-fold stage, the enriched area formed a finger-like extension from the ventral surface into the cytoplasm of the cell (Fig. 1D; n = 14 embryos).

**Figure 1.**
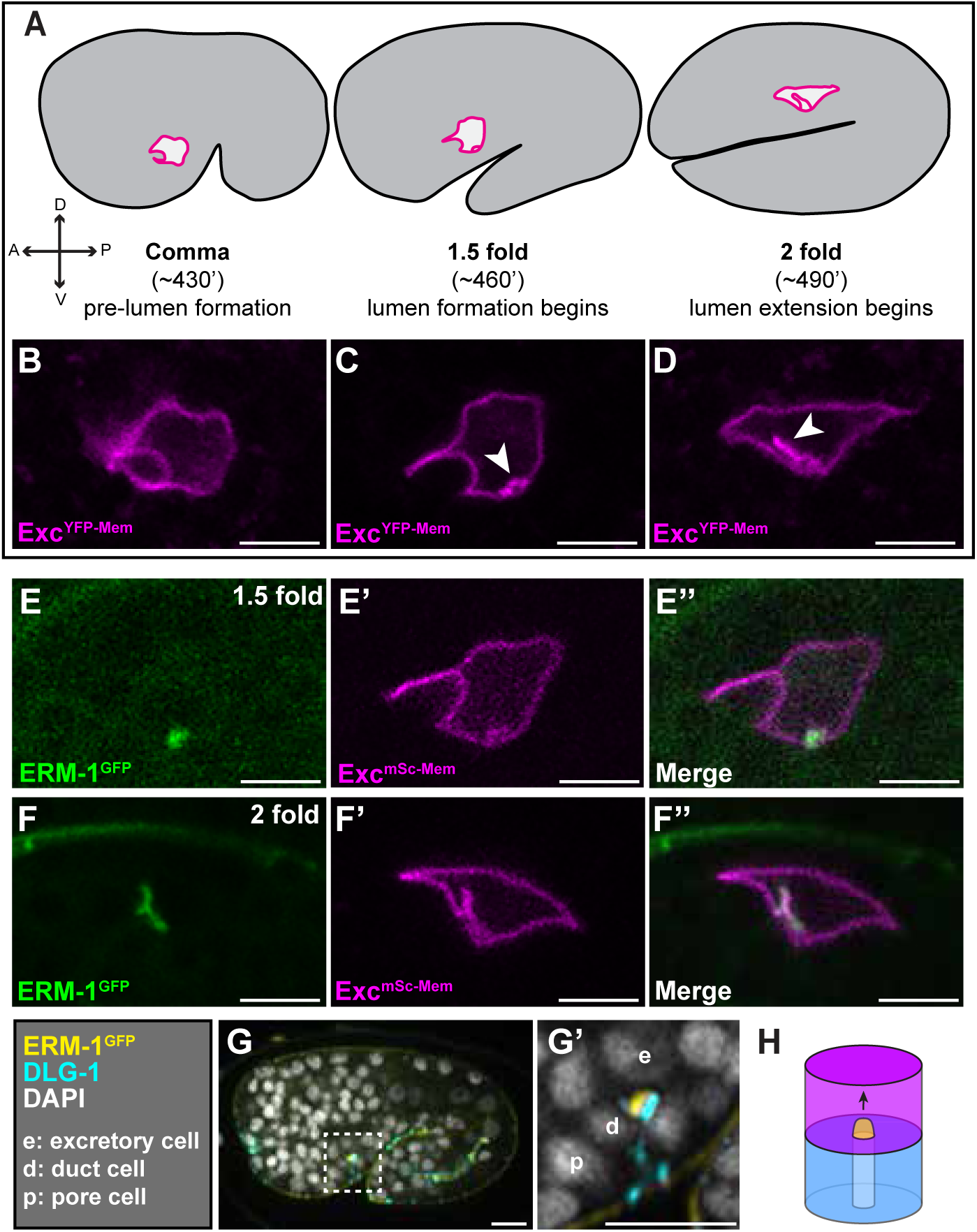
Lumen growth initiates at the plasma membrane adjacent to the duct cell. **(A)** Schematic of embryo and excretory cell morphology at comma, 1.5-fold and 2-fold stages from a lateral view; times listed are approximate minutes after the first embryonic cell division (20°C). **(B-D)** Expression of Exc^YFP-Mem^ (from *lin-3p::YFP::PH*) at comma stage (B), 1.5-fold stage (C), and 2-fold stage (D); arrowhead, nascent lumen forming at the ventral surface of the cell (C) and growing into the cytoplasm (D). **(E-F’’)** Co-localization of endogenously tagged ERM-1^GFP^ and Exc^mSc-Mem^ (from *lin-3p::mScarlet::PH*) at the excretory cell lumen in 1.5-fold (E-E’’) and 2-fold (F-F’’) embryos. **(G-G’)** Embryo stained for ERM-1^GFP^, junction marker DLG-1, and DNA dye DAPI. Nuclei of cells of the excretory system labeled as indicated. Inset of boxed region shown in G’. **(H)** Model of lumen initiation by apical invasion. Excretory cell is magenta, duct cell blue, and excretory lumen yellow. Scale bars, 5 µm.

We suspected that the ventral Exc^YFP-Mem^ enrichment marked the nascent excretory cell lumen. To test this hypothesis, we created an mScarlet version of the excretory cell plasma membrane marker (Exc^mSc-Mem^) and co-expressed it with endogenously tagged ERM-1^GFP^ (Ramalho et al., 2020). ERM-1 marks the excretory cell lumen in addition to apical surfaces of intestinal and gonadal epithelial cells (Gobel et al., 2004). The ventral membrane enrichment and finger-like extension marked by Exc^mSc-Mem^ colocalized completely with ERM-1^GFP^ in 15/15 1.5-fold and 5/5 2-fold embryos (Fig. 1E-F’’). In immunostained embryos, the initial ventral enrichment of ERM-1^GFP^ was invariably positioned adjacent to the duct cell contact (24/24), which we visualized by staining for the junction marker DLG-1 (Firestein and Rongo, 2001) (Fig. 1G; inset in G’). We conclude that the ventral membrane enrichment highlighted by our markers and ERM-1^GFP^ forms adjacent to the duct cell, extends inward into the excretory cell cytoplasm, and corresponds to the nascent excretory cell lumen (Fig. 1H). Given its narrow diameter, we were unable to resolve whether the lumen has a hollow center at this stage, although electron micrographs of fixed embryos indicate that this is the case (Berry et al., 2003; Stone et al., 2009). These findings strongly support an apical membrane invasion mechanism of lumen formation.

### Visualizing lumen branching in living embryos

Our efforts to use live imaging of Exc^YFP-Mem^ to watch the next steps in excretory cell lumenogenesis – left-right lumen growth and anterior-posterior splitting – were confounded by embryo twitching, which begins at the 2-fold stage. Twitching is powered by body wall muscle contractions, requires the muscle-specific myosin MYO-3, and is essential for the embryo to continue elongating beyond the 2-fold stage (Waterston, 1989; Williams and Waterston, 1994). *myo-3* mutant embryos are <u>p</u>aralyzed and <u>a</u>rrested at the two-fold stage (the <u>Pat</u> phenotype) but show signs of continued development, such as pharyngeal morphogenesis (Waterston, 1989), raising the possibility that *myo-3* mutant embryos could be used to visualize later stages of excretory cell lumenogenesis.

We were able to visualize continued development of the excretory cell in 3D timelapse movies of *myo-3* embryos expressing Exc^YFP-Mem^. Embryos were imaged from a left lateral perspective through the entire depth of the excretory cell. A projection of the complete image stack was rotated 90° along the anterior-posterior axis to present a dorsal view, which allows for left-right and anterior-posterior branching events to be visualized in one image. Because the right side of the cell (top of image panels) is deeper in the embryo and farther from the objective lens, fluorescence signals appear dimmer and less sharp than on the left side of the cell (bottom of image panels). At this stage of development, *lin-3p*-driven Exc^YFP-Mem^ was also expressed at a lower level in the adjacent excretory gland cells (Fig. 2A).

**Figure 2.**
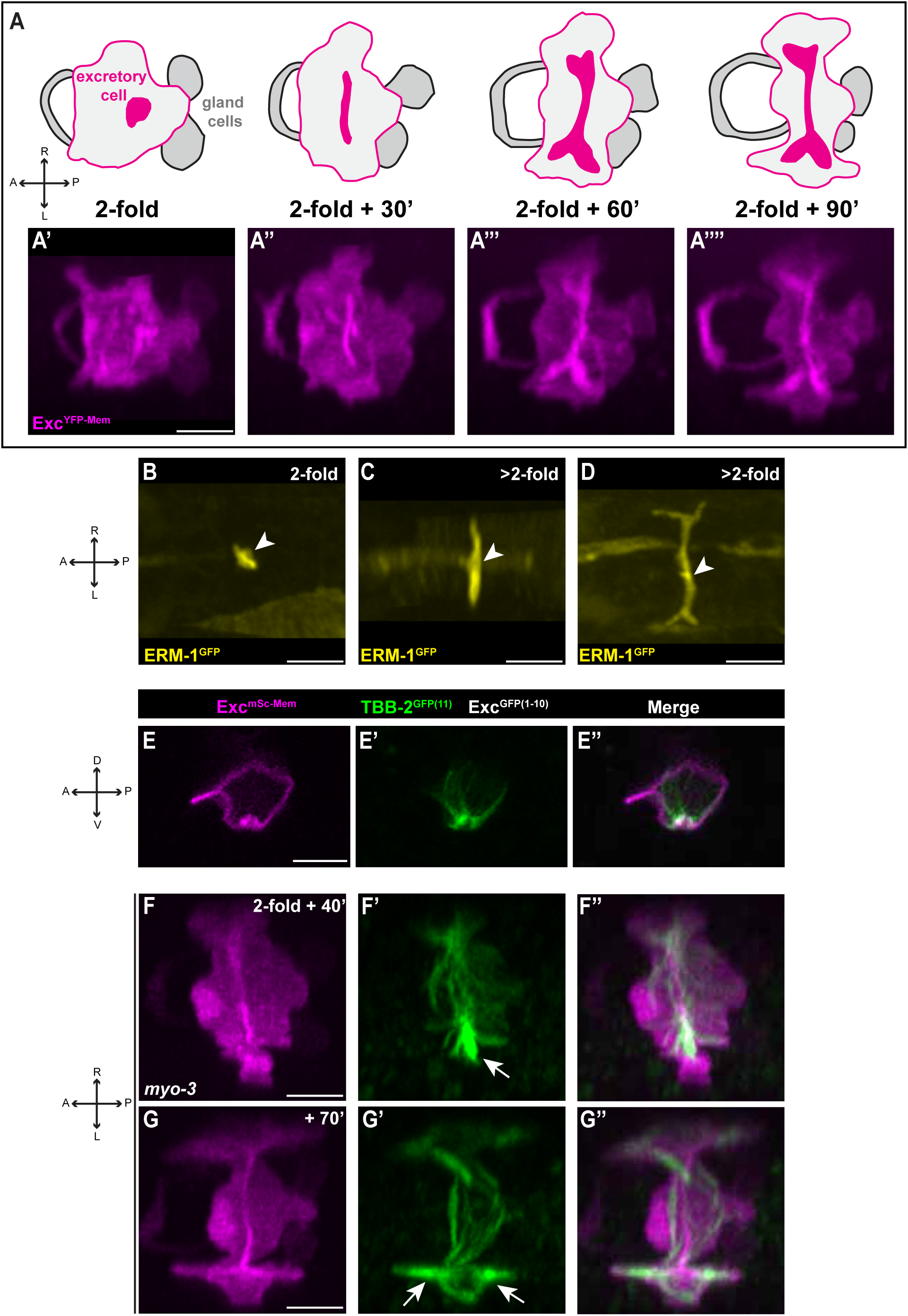
Excretory cell lumen branching and microtubule localization. **(A-A’’’’)** Dorsal view of excretory cell and lumen growth and branching; timelapse images from the 2-fold stage onward in *myo-3* mutant embryos. Exc^YFP-Mem^ labels the cell membrane and lumen of the excretory cell and is also expressed at lower levels in the gland cells (gray shading in schematic). **(B-D)** Excretory cell lumen in embryos stained for ERM-1^GFP^ (anti-GFP) at stages like those shown in live embryos in (A’-A’’’). **(E-G’’)** Exc^mSc-Mem^ and endogenous MTs visualized live by split GFP labeling of TBB-2/ß-tubulin. Images in E-E’’ are from a lateral view; images in F-G’’ are timelapse stills from a dorsal view. Arrows indicate MT concentrations near the end of the growing lumen. Scale bars, 5 µm.

In timelapse movies of 15 *myo-3* mutant embryos started at the initiation of the 2-fold stage, we observed that the excretory cell began to extend along its left-right axis (Fig. 2A’’). Simultaneously, the nascent lumen split and began to grow along the left-right axis, mirroring the excretory cell extensions and forming what will become the crossbar of the H shape. After the left and right cellular extensions reached the left and right lateral edges of the embryo, the cell and the trailing lumen branched to extend along the anterior-posterior axis in both directions (Fig. 2A’’’,A’’’’). This second branching event created the four arms of the H. Although we cannot exclude the possibility that the 2-fold arrest or lack of twitching in *myo-3* mutants affects excretory cell branching and early growth in subtle ways, this sequence of events matches those of fixed 2-fold and older wild-type embryos expressing ERM-1^GFP^ and immunostained for GFP to mark the lumen (Fig. 2B-D, arrowheads; n = 43 embryos).

### Microtubules concentrate near the ends of extending lumen branches

Previous studies have shown a role for microtubule (MT) regulation in excretory cell lumen development in larvae (Shaye and Greenwald, 2015). MTs visualized indirectly using the MT-binding domain of Ensconsin show a filamentous localization along the length of the larval excretory cell and are enriched at the tips of the excretory cell extensions. To examine the dynamics of endogenous excretory cell MTs as the lumen first develops and branches in embryos, we utilized a tissue-specific split GFP approach (Noma et al., 2017). For these experiments, we endogenously tagged *tbb-2/β-tubulin* with sequences encoding GFP11 (the 11^th^ β-strand of GFP), expressed the complementing GFP1-10 fragment from the *lin-3* promoter, and visualized MTs in timelapse movies of 18 *myo-3* mutant embryos. At the initial stages of lumen appearance at the ventral surface, MTs accumulated at this site and radiated inward into the cell body (Fig. 2E-E’’). After the lumen branched along the left-right axis, MTs became concentrated near the tips of the growing lumen and were present at lower density in the remainder of the cell and along already-formed regions of lumen (Fig. 2F-F’’). This pattern continued after the anterior-posterior lumen split, with each of the four tips of extending lumen showing a concentrated MT accumulation (Fig. 2G-G’’). We conclude that in the embryonic excretory cell, MTs first enrich at the site of lumen initiation, then as the lumen extends, MTs concentrate near the growing lumen tips, with the MT concentration close to but not reaching the leading edge of the excretory cell extensions.

### INA-1/PAT-3 integrin is required for excretory cell morphogenesis

We searched for regulators of excretory cell lumen patterning by screening for EMS-induced mutants with defects in excretory cell morphogenesis. To facilitate mutant identification, we created a strain expressing endogenously tagged (pHluorin) lumen marker VHA-5 (Kolotuev et al., 2013) and screened F_2_ progeny of mutagenized worms using a fluorescence dissection scope. Three lethal mutants (*xn182*, *xn190*, and *xn191*) caused an almost complete failure of excretory cell extension. Utilizing polymorphism mapping and whole genome sequencing (see Methods), we identified *xn182* and *xn190* as missense mutations within the *ina-1* gene (Fig. 3A,B). To confirm that *ina-1* disruption was the cause of the excretory cell defects, we introduced the *xn182* missense mutation into a non-mutagenized background using CRISPR/Cas9, generating *xn219*. Whereas wild-type L1 excretory canals extended over half the distance from the cell body to the posterior of the worm (Fig. 3C,F), *ina-1(xn219)* L1 larvae had severely truncated excretory cells, similar in severity to those of presumptive null mutant *ina-1(gm86)* (Fig. 3D-F).

**Figure 3.**
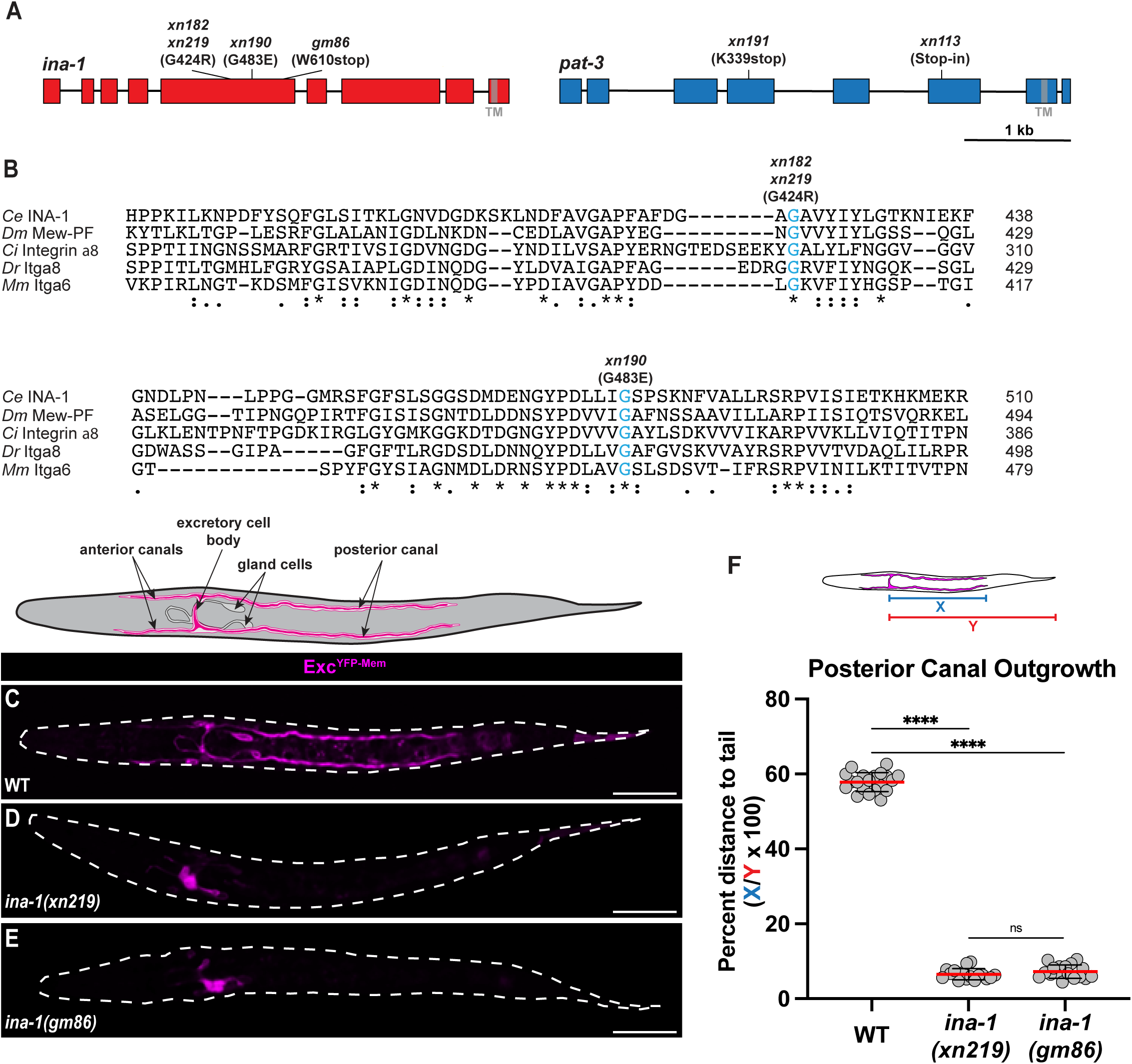
Mutations in integrin prevent excretory canal outgrowth. **(A)** Schematic of *ina-1* and *pat-3* genes indicating mutations; exons (boxes), introns (lines), and region encoding the transmembrane domain (‘TM’; shaded gray) are shown. *ina-1(xn219)* is a CRISPR recreation of the *ina-1(xn182)* allele. The *ina-1(gm86)* allele contains a missense mutation (A565T) in addition to the annotated nonsense mutation (Baum and Garriga, 1997). The *pat-3(xn113)* allele contains a STOP-IN cassette inserted at the indicated position; the STOP-IN cassette causes a frameshift and premature stop, and was created by editing the *pat-3(xn109[pat-3::zf1::gfp])* allele (McIntyre and Nance, 2023). **(B)** Portion of the INA-1 ß propeller domain containing mutated residues, with orthologues from fruit fly *Drosophila melanogaster* (*Dm*), sea squirt *Ciona intestinalis* (*Ci*), zebrafish *Danio rerio* (*Dr*), and house mouse *Mus musculus* (*Mm*). Symbols below indicate degree of amino acid conservation (asterisk – identical; colon – strongly similar; period – weakly similar). Conserved glycine residues mutated in *xn182* and *xn190* are shaded blue. **(C-E)** Exc^YFP-Mem^ in wild type (‘WT’), *ina-1(xn219)* and *ina-1(gm86)* L1s (larval body outlined in dashed white); schematic above shows relevant anatomical features. **(F)** Posterior canal outgrowth, calculated as percent distance to tail as schematized, in L1 larvae of the indicated genotypes. Circles: individual worms, red lines: mean, black lines: error bars (S.D.). ****p < 0.0001, ns = not significant (p > 0.05). Scale bars, 25 µm.

*ina-1* is one of two ⍺-integrin genes in *C. elegans* (Baum and Garriga, 1997). Integrins are heterodimeric basement membrane receptors composed of single ⍺ and β subunits, each of which contains extracellular, transmembrane, and cytoplasmic domains (Kanchanawong and Calderwood, 2023). Both *xn182* (G424R) and *xn190* (G483E) convert highly conserved glycine residues within the extracellular ß-propeller domain, which is involved in ligand binding and ⍺/β dimer formation, to charged residues (Fig. 3B). INA-1, whose closest mammalian ortholog is Itga6, is homologous to the class of ⍺-integrins that binds to the basement membrane protein laminin. The other *C. elegans* ⍺-integrin, PAT-2, belongs to the class that binds RGD peptides (Hynes and Zhao, 2000).

*C. elegans* has only a single β-integrin gene, *pat-3* (Baum and Garriga, 1997; Gettner et al., 1995). The *xn191* mutant introduced a nonsense mutation within *pat-3* prior to the transmembrane domain. Like *myo-3* mutants and previously characterized *pat-3* null mutants (Williams and Waterston, 1994), *pat-3(xn191)* mutants arrested during embryogenesis with a Pat phenotype. As we detail below, *pat-3* mutant embryos have severe excretory cell morphogenesis defects, like *ina-1* mutants. Together, these findings indicate that INA-1/PAT-3 heterodimers regulate excretory cell morphogenesis.

### Integrin is required for anterior-posterior branching of excretory cell extensions

We used live imaging of Exc^YFP-Mem^ to determine which step of excretory cell morphogenesis and lumen formation is affected by mutations in integrin. In both *ina-1* (18/18) and *pat-3* (23/23) mutants, the initial formation of the nascent lumen at the ventral surface of the excretory cell occurs normally (Fig. 4A-C). Because *pat-3* mutants show the same Pat phenotype as *myo-3* mutants, we were able to examine subsequent stages of lumenogenesis in timelapse movies. The growth of the excretory cell and the splitting of its lumen along the left-right axis occurred successfully in *pat-3* mutants (Fig. 4D; compare to Fig. 2A’’). However, in contrast to *myo-3* mutants (examined at 2-fold plus 90’), the anterior-posterior branching of the excretory cell and lumen failed to occur in 16/18 mutants at the same developmental stage (Fig. 4E; compare to Fig. 2A’’’’), and the small fraction that did split had severely truncated projections that failed to continue extending. Even though the cell failed to extend anterior-posterior projections, the lumen appeared to continue to expand within the cell and became irregular in shape and width, suggesting that lumenal membrane continues to be added in *pat-3* mutants even though the cell itself fails to extend. (Fig. 4 D-D’’). Anterior-posterior excretory cell branching occurred successfully in 10/10 *pat-2/⍺-integrin* mutant embryos (Fig. S1), which also arrest as paralyzed two-fold embryos (Williams and Waterston, 1994), consistent with our interpretation that this stage of excretory cell morphogenesis is mediated by INA-1/PAT-3 heterodimers.

**Figure 4.**
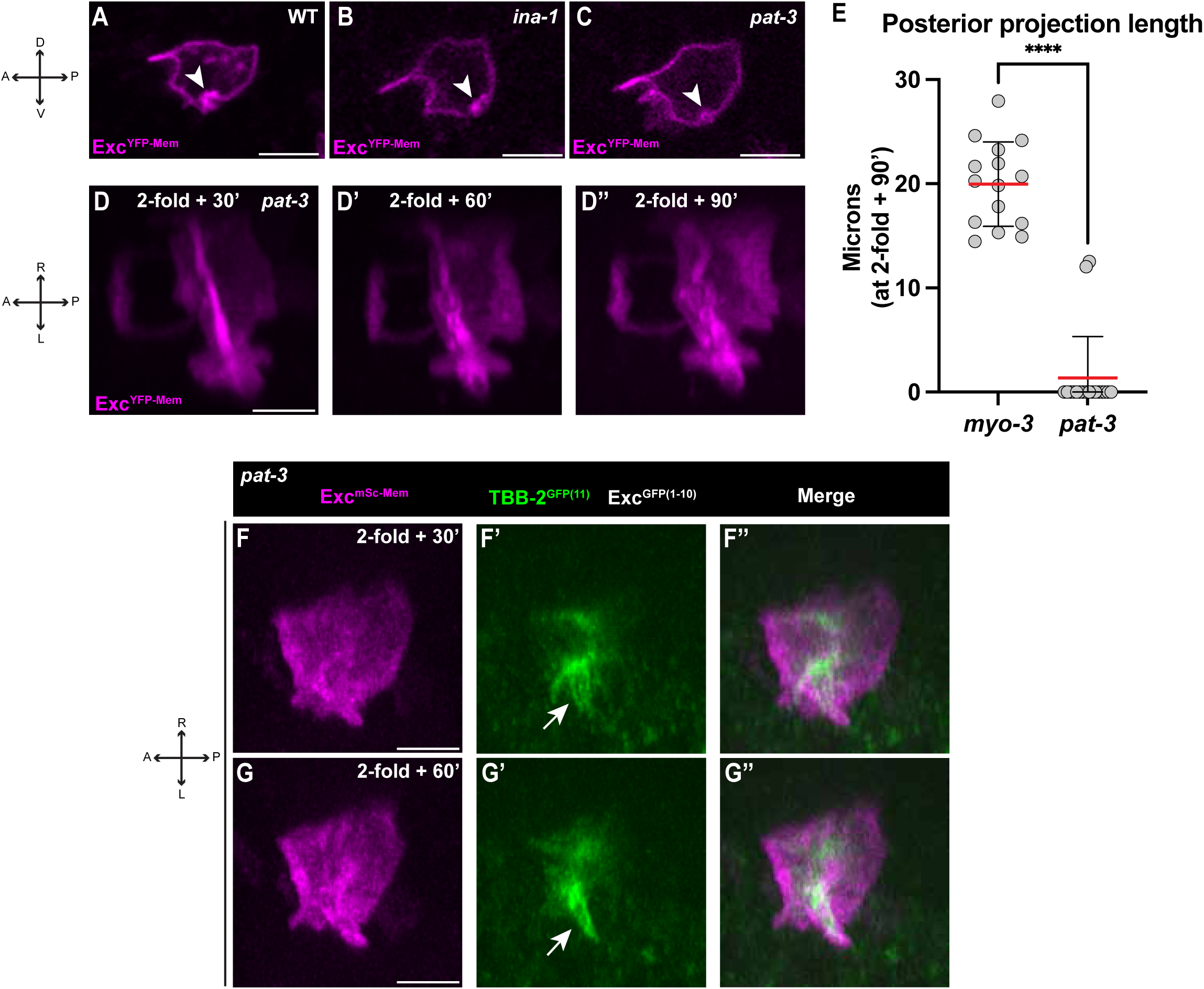
Lumen formation and microtubule localization in integrin mutants. **(A-C)** Exc^YFP-Mem^ in 1.5-fold WT, *ina-1(xn219),* and *pat-3(xn113)* embryos; arrowheads indicate the nascent lumen. **(D-D’’)** Timelapse stills of Exc^YFP-Mem^ in *pat-3(xn113)* embryos at the indicated times after 2-fold arrest. **(E)** Posterior excretory cell projection length at 2-fold + 90’ of the indicated genotypes. Circles: individual worms, red lines: mean, black lines: error bars (S.D.). ****p<0.0001. **(F-G’’)** Timelapse stills of Exc^mSc-Mem^ and endogenous MTs in *pat-3(xn113)* embryos visualized by split GFP labeling of TBB-2/ß-tubulin. Arrows indicate the concentration of MTs at the end of the growing lumen. Scale bars, 5 µm.

The pattern of MTs in *pat-3* mutant embryos was also consistent with a defect specifically in the anterior-posterior branching of the excretory cell lumen: microtubules concentrated normally around the tips of the left-right branches in *pat-3* embryos before becoming disorganized at later stages (Fig. 4F-G’’). Based on these observations, we conclude that integrin is dispensable for the initial formation of the excretory cell lumen and for lumen left-right branching but is needed for the cell and lumen to form anterior-posterior branches. At a gross level, MT organization occurs normally in *pat-3* embryos until the stage when anterior-posterior branching should have occurred.

### INA-1 functions cell autonomously to promote excretory cell morphogenesis

The excretory cell interacts with several other cell types as it extends and branches, raising the question of whether integrin functions cell autonomously within the excretory cell or non-autonomously in other cells. Given the numerous potential sites of action, we carried out an unbiased genetic mosaic analysis to define where *ina-1* function is needed. For this approach, we used an extrachromosomal array *(xnEx562)* containing *ina-1(+)* and the *sur-5::gfp* co-injection marker, whose product marks the nuclei of most cells, to rescue *ina-1(xn219)* mutants. At each embryonic cell division, extrachromosomal arrays are typically inherited by both daughter cells. However, in rare cases, one daughter cell does not inherit the array; such animals are genetic mosaics. For *ina-1; xnEx562* animals, a cell that does not inherit the array and each of its descendants lack *ina-1* gene function and can be recognized as such by lack of SUR-5^GFP^ in their nuclei. Most *ina-1; xnEx562* animals showed normal excretory cell morphogenesis, which we scored in late larvae and adults using a transgene expressing excretory-specific cytoplasmic mCherry (*pgp-12p::mCherry*). We searched for rare mosaic animals with defective excretory cells that failed to undergo morphogenesis. In 28/28 mosaics with the mutant phenotype, SUR-5^GFP^ was absent from the excretory cell nucleus (but present in many other nuclei); by contrast, in 83/83 worms with normal excretory cell branching, SUR-5^GFP^ was present in the excretory cell (Fig. 5A-B’). Based on these observations, we infer that all mosaic animals with excretory cell morphogenesis phenotypes lost the *xnEx562* rescuing array at some point within the lineage that gives rise to the excretory cell. These findings also demonstrate that excretory cell canal extension is not needed for larval development, and that the L1 lethality observed in *ina-1* null mutants (Baum and Garriga, 1997) arises from an unknown, essential *ina-1* function in another tissue(s).

**Figure 5.**
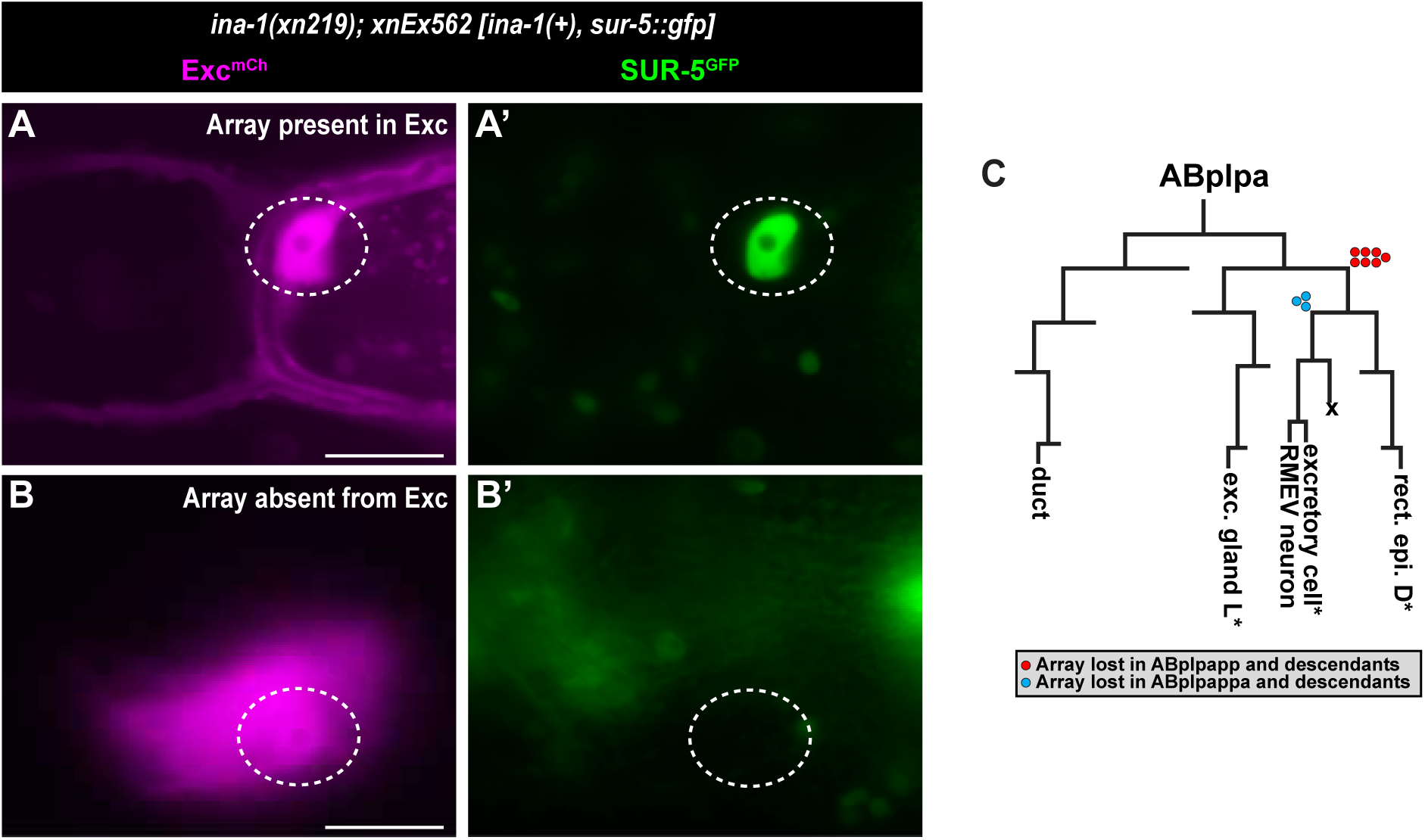
*ina-1* mosaic analysis. **(A-B’)** Expression of SUR-5^GFP^ and Exc^mCh^ (from *pgp-12p::mCherry*) in adult *ina-1* mutants carrying the *xnEx562 [ina-1(+), sur-5::gfp]* rescuing extrachromosomal array. (A) Worm with properly formed excretory cell; SUR-5^GFP^ expression in the nucleus (dashed circle) indicates the array is present in the excretory cell. (B) Worm with defective excretory cell outgrowth; lack of SUR-5^GFP^ indicates the rescuing array was lost at some point in the lineage giving rise to the excretory cell. **(C)** ABplpa lineage, which gives rise to the excretory cell. Colored circles: mosaic animals with defective excretory cells that resulted from loss of the array at the indicated step in the lineage. Cells marked with asterisk were scored for SUR-5^GFP^ expression to infer the loss-point; red circles, array loss at ABplpapp; blue circles: loss at ABplpappa or later. Scale bars, 5 µm.

The excretory cell descends from the ABplpa blastomere, which also gives rise to the duct cell and the left excretory gland cell (Fig. 5C). To determine if the *xnEx562* array was required in the excretory cell to rescue the excretory morphogenesis phenotype of *ina-1* mutants, we searched for animals that were mosaic within the ABplpa lineage. To find ABplpa mosaics, we scored *ina-1; xnEx562* animals with defective excretory cell morphogenesis for SUR-5^GFP^ expression in three nuclei within the ABplpa lineage (excretory cell, rectal epithelial cell D, left excretory gland cell; asterisks, Fig. 5C). We were able to identify ten ABplpa mosaics and inferred the array loss point by the combination of these three cells that expressed SUR-5^GFP^ (Fig. 5C; colored dots are inferred loss points); in three of these mosaics (blue dots), the array loss occurred late in the lineage, affecting only the excretory cell and potentially its sister, the RMEV motor neuron (which is located in the nerve ring and whose nucleus we were unable to score). Altogether, these results strongly support a cell autonomous function for *ina-1* in excretory cell branching and extension. Consistent with this interpretation, *ina-1* mRNA detected by single-cell mRNA sequencing is present in the excretory cell at its birth and throughout the embryonic stages when excretory morphogenesis occurs (Packer et al., 2019).

### Integrin is required for the excretory cell to cross the epidermal basement membrane

Basement membrane (BM) first assembles during mid-embryogenesis on the basal surfaces of internal organs (e.g. the pharynx) and surface epidermal cells (Huang et al., 2003; Kao et al., 2006; Keeley et al., 2020). Whereas the body of the excretory cell is positioned in the interior of the embryo adjacent to the pharynx, the anterior and posterior excretory canals migrate between the epidermis and the epidermal BM (Hedgecock et al., 1990; Nelson et al., 1983; White et al., 1986). This anatomical organization raises the question of how the excretory canals traverse from one side of the epidermal basement membrane to the other as they extend from the cell body.

Given the role of integrins in cell-basement membrane interactions, we investigated the relationship between the excretory cell, epidermal BM, and integrin function. BM formation begins when laminin assembles into a scaffold on cell surfaces. Laminin is a heterotrimer composed of α, β, and γ subunits (Yurchenco and Kulczyk, 2024); *C. elegans* contains two α subunit genes (*epi-1* and *lam-3*) and unique β (*lam-1*) and γ (*lam-2*) subunit genes (Huang et al., 2003; Kao et al., 2006). To examine BM and the excretory cell simultaneously, we imaged L1 larvae co-expressing excretory-specific cytoplasmic mCherry (*pgp-12p::mCherry*) and endogenously tagged LAM-2 fused to mNeonGreen (LAM-2^mNG^), which marks all BMs (Keeley et al., 2020). In 22/22 canal branches from 11 wild-type animals, the excretory cell canals passed through a prominent gap in the epidermal BM (Fig. 6A-A’’, arrow). We quantified the presence of the gap and its position by plotting normalized intensity measurements of LAM-2^mNG^ along an 8 µm line centered where the excretory cell splits to form anterior and posterior canals, which revealed a several micron gap in LAM-2^mNG^ centered on the excretory cell branching point (Fig. 6C, average values; Fig. 6E, individual traces in white). Remarkably, in *ina-1* mutant L1s, the epidermal BM lacked a gap at this position (0/20 canal branches from 10 worms contained a BM gap) (Fig. 6B; Fig. 6D, average values; Fig. 6E, individual traces in black). We conclude that the excretory cell extensions normally cross the BM and *ina-1* is required for it to do so.

**Figure 6.**
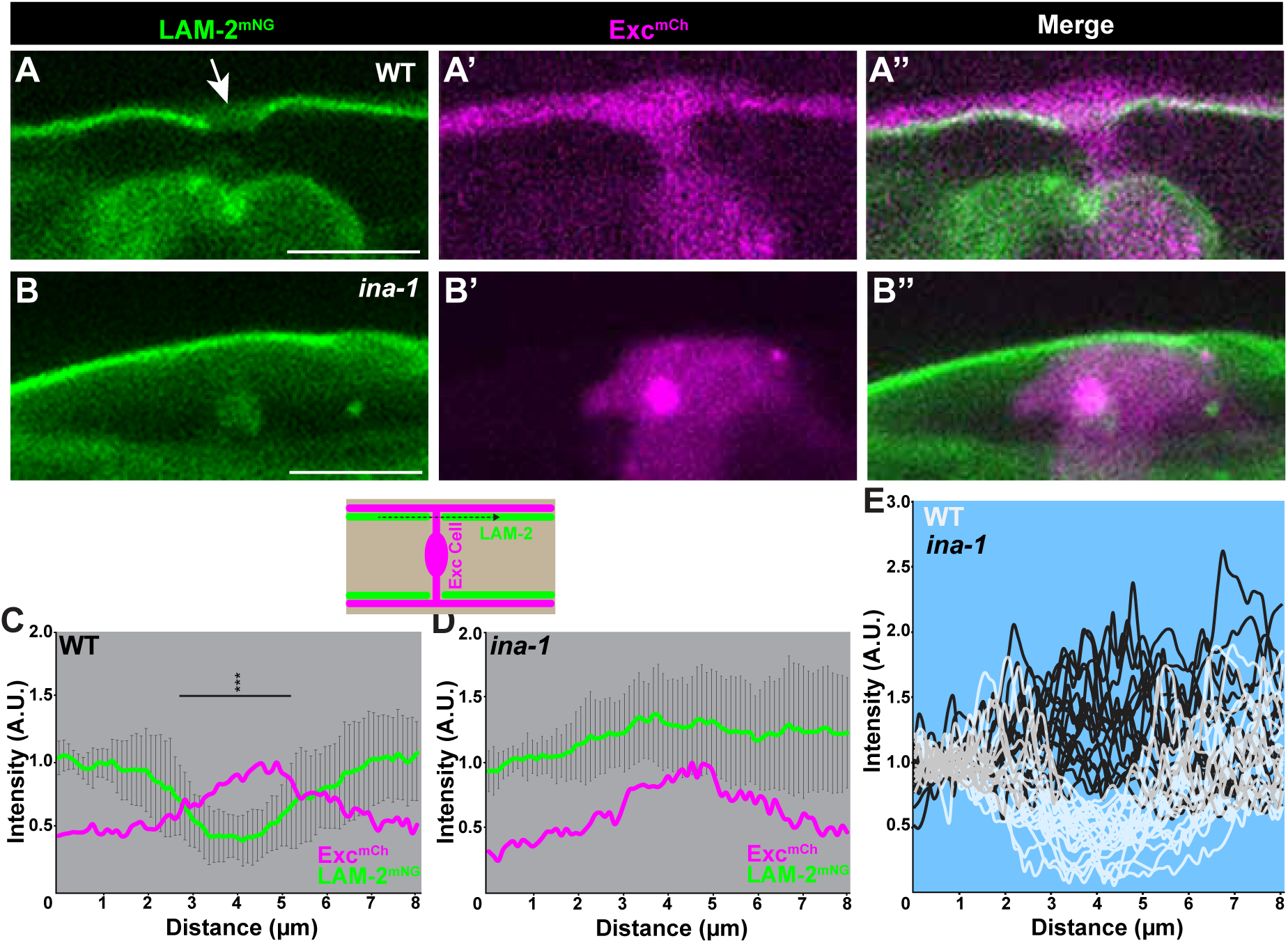
Excretory cell integrin-dependent basement membrane crossing. **(A-B’’)** LAM-2^mNG^ and cytoplasmic Exc^mCh^ in wild-type (A-A’’) or *ina-1(xn219)* (B-B’’) L1; arrow, basement membrane gap in wild-type. **(C-D)** Normalized LAM-2^mNG^ and Exc^mCh^ intensities in wild-type (C) and *ina-1(xn219)* (D); schematic above shows region of measurements taken along an 8µm span of basement membrane centered on the excretory cell. Statistical comparisons were between the most posterior (last) LAM-2^mNG^ measurement and all others using a t-test with Bonferroni correction for multiple comparisons. Error bars, S.D. ***p < 0.001. **(E)** Individual data plots of normalized LAM-2^mNG^ intensities that were used for averages shown in C-D. Black lines, wild-type; white lines, *ina-1* (overlap of lines is gray). Scale bars, 5µm.

## DISCUSSION

Numerous studies in *C. elegans* larvae have revealed mechanisms of excretory cell lumen growth and maintenance by cytoskeletal proteins and trafficking pathways (Shaye and Soto, 2021; Sundaram and Buechner, 2016). However, the lumen first begins to form and acquires its H-shaped pattern during embryogenesis. To date, live imaging has not been used to examine early excretory cell lumenogenesis because embryo movements prevent timelapse imaging during this stage. Here, we show that excretory cell development continues in genetically paralyzed animals, and we take advantage of these mutants to examine early stages of excretory cell lumenogenesis using live imaging. Our findings provide new insights into two essential aspects of excretory cell tubulogenesis: how the intracellular lumen first initiates, and how its stereotypical pattern of branching is regulated by integrins and basement membrane.

### Lumen initiation

Previous studies of lumen initiation examined fixed embryos using transmission electron microscopy (TEM) and arrived at different models of how lumen formation begins. One TEM analysis of serial sections noted a large vacuole within the excretory cell cytoplasm just prior to detecting an elongated lumen. Based on this finding, the authors proposed that cytoplasmic vacuole coalescence initiated the lumen, which then expanded and connected to the duct cell lumen (Berry et al., 2003). A subsequent examination of existing TEM sections identified an embryo showing an apparent nascent lumen extending from the duct cell connection into the excretory cell cytoplasm, leading the authors to suggest that the lumen initiated at this site (Stone et al., 2009). Our analysis of lumen initiation in living embryos provides strong evidence for this second model. First, the initial evidence of lumen formation that we detected was invariably at the site of contact with the duct cell and began as a small membrane thickening that co-localized with the ERM-1^GFP^ lumenal marker. This thickening subsequently expanded into the excretory cell cytoplasm to form a finger-like invasion. The PH_PLC∂1_ membrane marker, which is first expressed well before lumenogenesis begins and enriches both at the plasma membrane and nascent lumen, never concentrated on a vacuole-like endomembrane. While it is possible that such a vacuole is composed of lipids not efficiently recognized by the PH_PLC∂1_ domain until the vacuole fuses with the plasma membrane, the very small size of the initial lumen that we detected, and its invariant position at the excretory-duct interface, seem at odds with this model.

Previous studies have shown that lumen extension is driven by vesicle trafficking pathways, which are thought to provide the additional membrane needed for lumenogenesis. For example, components of the exocyst vesicle-tethering complex are recruited to the lumenal membrane by PAR-6, and degron-mediated depletion of the exocyst or PAR-6 results in a severely truncated excretory cell lumen (Abrams and Nance, 2021; Armenti et al., 2014). Therefore, local activation or recruitment of the exocyst, or other trafficking pathways, could be one mechanism triggering apical membrane invasion. In future studies, it will be important to identify the signal that initiates apical invasion and determine whether this is supplied by the duct cell or occurs as a programmed step in excretory cell differentiation.

The initiation of intracellular lumenogenesis has been examined *in vivo* in *Drosophila* terminal tracheal cells. Like the excretory cell, terminal tracheal cells extend a blind-ended intracellular lumen from the site of contact with the existing tracheal lumen (Gervais and Casanova, 2010). The terminal tracheal cell lumen initiates from a small ring-like apical patch, which then extends into the cell. Also similar to the excretory cell, microtubules initially emanate from the apical patch, then concentrate near the tip of the extending lumen (Gervais and Casanova, 2010). Apical membrane invasion has been observed in other examples of intracellular tubes, including lumenizing endothelial cells during vessel anastomosis in zebrafish (Lenard et al., 2013) and in notochord formation in ascidian embryos (Dong et al., 2009). These similarities suggest the presence of a conserved molecular mechanism of intracellular lumen formation that operates across a wide variety of cell types.

### Mechanisms of lumen branching

Unlike *Drosophila* terminal tracheal cells, whose branching pattern varies between embryos and even between homologous cells on contralateral sides of the embryo (Gavrilchenko et al., 2024), the excretory cell branches in a stereotypical pattern to form its H-shaped canals (Nelson et al., 1983). The molecular mechanisms that guide this invariant branching pattern are largely unknown. Our screen for excretory cell lumen patterning mutants identified integrins (*ina-1* and *pat-3*) as key regulators of excretory cell lumen branching, with a cell-autonomous requirement to transition from left-right to anterior-posterior lumen extension.

The excretory cell extensions pass through the epidermal BM at the site where the anterior-posterior branching occurs, and we found that this process fails in *ina-1* mutants. Failure of the excretory cell to cross the epidermal basement membrane could be explained by several potential defects. One possibility is that the cell branches prior to epidermal BM formation and the BM is deposited around the already-branched cell. We consider this very unlikely since the epidermal BM assembles prior to embryo elongation (Keeley et al., 2020), well before excretory cell branching begins. A second possibility is that another cell requires *ina-1* to breach the epidermal BM and the excretory cell migrates through this BM hole. While we cannot completely exclude this possibility, our finding that *ina-1* functions cell autonomously within the excretory cell for anterior-posterior branching argues strongly against it. A final possibility is that the excretory cell itself breaches the epidermal BM and requires INA-1/PAT-3 to do so. We favor this model because our data suggest a cell-autonomous function of *ina-1* in the excretory cell.

We envision several possible mechanisms for how INA-1/PAT-3 integrin promotes lumen branching. One possibility is that integrin function is needed solely for basement membrane breaching, which is a pre-requisite for lumen branching. For example, signals that promote bidirectional migration of the excretory cell extensions (towards both the anterior and posterior) could be inaccessible to the excretory cell if it does not cross the basement membrane. There is precedent for this function of integrin in another cell type that undergoes a programmed breaching of basement membrane – the *C. elegans* anchor cell (AC). Analogous to the excretory cell, the AC requires cell autonomous *ina-1* function to penetrate the uterine basement membrane (Hagedorn et al., 2009). *ina-1* also regulates several cell migrations that occur during *C. elegans* development, such as the posteriorly directed migration of the CAN neurons (Baum and Garriga, 1997), and thus could have a subsequent function in the anterior-posterior migration of the excretory cell extensions. Addressing this possibility will require new genetic methods to acutely inhibit integrin function just after the excretory cell successfully crosses the epidermal basement membrane.

Integrin function has also been shown to regulate *Drosophila* terminal tracheal cell patterning, but in this context appears to have a distinct cellular function. In tracheal cells, the ß-integrin Myospheroid functions with two different α-integrins and the downstream effector talin to promote the maintenance of tracheal cell branches; when integrin signaling activity is blocked, terminal tracheal cell lumenogenesis occurs normally but the branches eventually deteriorate, perhaps due to defective adhesion with the underlying basement membrane (Levi et al., 2006). Given the early block in excretory cell morphogenesis in *C. elegans* integrin mutants, it is not clear whether integrin function might also be required for continued adhesion of the excretory cell canals to the epidermal basement membrane on which they reside. Despite these differences in integrin mutant phenotype, it is possible that the molecular function of integrin is similar in both cases – cell adhesion to the BM could be required for excretory cell BM invasion in *C. elegans* and maintenance of tracheal tubes in *Drosophila*.

## METHODS

### Worm culture and strains

All strains were maintained at room temperature and raised on NGM plates seeded with *Escherichia coli* strain OP50 (Brenner, 1974). Strains created or used in this study are listed in Table S1.

### Mutant screen

To mark the excretory cell, we created an endogenously tagged *vha-5(xn93[vha-5::pHluorin])* strain. Consistent with *vha-5* transgene expression studies (Liegeois et al., 2006), *vha-5(xn93[vha-5::pHluorin])* worms show bright green fluorescence in the excretory cell as well as the epidermis. Synchronized L4 *vha-5(xn93)* worms were mutagenized with 50 mM EMS (Brenner, 1974). The following day, eggs were collected from mutagenized worms and allowed to hatch overnight in M9 buffer before plating as L1 larvae split onto 60 seeded NGM agar plates. Plates were placed at staggered temperatures to allow for screening on three consecutive days. From each set of 20 plates, 200 F_1_ L4 larvae were singled to individual plates and allowed to self for one generation (in later rounds of screening, three F_1_ were placed on each plate). F_2_ were screened for truncated excretory cell canals using a Leica M165C fluorescence dissection scope. After screening F_2_ broods from 7200 F_1_, three mutants were identified that blocked excretory cell outgrowth completely (*xn182, xn190, xn191*). All three mutations were homozygous lethal and were recovered from heterozygous siblings.

To identify candidate causal mutations, each mutant was backcrossed to the unmutagenized *vha-5(xn93[vha-5::pHluorin])* strain and homozygous mutant or non-mutant progeny were isolated in the F_2_ as described (Joseph et al., 2018). Because mutants were lethal, ∼200 mutant embryos (*xn191*) or L1 (*xn182, xn190*) were picked into M9 for DNA prep as described (Smith et al., 2016). A standard phenol chloroform extraction was performed on ∼50 pooled non-mutant plates confirmed to be homozygous wild-type by the lack of phenotypes in the F_3_. Libraries were then constructed using the Kapa HyperPrep kit (Roche) and paired-end sequences were obtained using an Illumina NovaSeq 6000. Candidate causal mutations were (1) those present in the mutant pool and absent in the non-mutant pool, and (2) predicted to alter protein sequence (missense, nonsense, and splice donor or acceptor mutants) (Joseph et al., 2018). Because two candidate causal mutations mapped to *ina-1*, we introduced the *xn182* mutation into an otherwise wild-type background using CRISPR/Cas9 [creating *ina-1(xn219)*], which recapitulated the mutant phenotype. For *pat-3*, excretory phenotypes were confirmed in the *pat-3(xn113)* null allele (McIntyre and Nance, 2023), which like *pat-3(xn191)* creates an early stop prior to the transmembrane domain. The phenotype of *pat-3(xn191)* was later characterized when we introduced the mutation into a line expressing *2xGFP11::tbb-2* but that was otherwise wild type.

### Mosaic analysis

*ina-1(xn219)/qC1* worms expressing the *xnSi37* [*pgp-12p::mCherry*] excretory cell cytoplasmic marker were injected with genomic DNA containing the *ina-1* gene (fosmid WRM068dG03, 20 ng/µl) and *sur-5::gfp* co-transformation marker (plasmid pTG96, 80 ng/µl) (Yochem et al., 1998) to produce a viable *ina-1(xn219); xnSi37; xnEx562 [ina-1(+), sur-5::gfp]* strain. Adults were scored under a fluorescence dissection scope to identify mosaics with defective excretory cell morphogenesis. Mosaics were transferred to agar pads and imaged on a Zeiss AxioImager A.2 wide-field epifluorescence microscope with 40x 1.3 NA lens. DIC, GFP and mCherry stacks through the animal were taken in the head region, central region, and tail region. Mosaics oriented with the left side facing the coverslip were scored for SUR-5^GFP^ in the excretory cell, excretory gland left, and rectal epithelial cell D (Yochem, 2006); mosaics with expression in some but not all of these cells were inferred to have lost the rescuing array in descendants of ABplpa (Sulston et al., 1983). Separately, adult animals were scored in bulk on agar pads to correlate successful excretory cell branching with SUR-5^GFP^ expression in the excretory cell nucleus.

### Transgene construction

All transgenes were constructed using Gibson assembly (Gibson et al., 2008) and cloned into MosSCI vector pCFJ151 (Frokjaer-Jensen et al., 2008), which contains the *Cbunc-119(+)* co-transformation marker, homology arms flanking the *ttTi5605* Mos insertion, and the *unc-54* 3’ UTR. The *lin-3* promoter fragment expresses in the embryonic excretory cell and excretory gland cells, as well as the larval anchor cell (Abdus-Saboor et al., 2011). The *pgp-12* promoter expresses in the larval and adult excretory cell (Zhao et al., 2005). *PH* encodes the PIP2-binding PH domain from rat PLC81, which targets fusion proteins to the plasma membrane (Audhya et al., 2005). *exc* encodes the apical targeting region of EXC-4 (Berry et al., 2003), but the fusion proteins we created using the *lin-3* promoter and *exc* show cytoplasmic rather than apical localization and were used as a marker of the cytoplasm. *GFP1-10* encodes the first 10 ß-strands of GFP (Noma et al., 2017). Relevant promoter and coding sequences of plasmids used in this study are provided in the Supplemental Material.

### Transgenesis and integration

Microinjections to create extrachromosomal arrays were performed as described (Mello et al., 1991). pLM006 and pLM007 as well as pLM008 and pLM009 were injected in pairs at 30 ng/µl into *unc-119(ed3)* young adult worms. pLM014 and pLM006 were injected as a pair into *tbb-2(xn226: 2xgfp11::tbb-2)* worms in both *myo-3(st386)/tmC12 and pat-3(xn234)/qC1* backgrounds with dominant Rol plasmid pRF4 as a co-injection marker at 100 ng/µl.

Extrachromosomal arrays containing pLM006 and pLM007, or pLM008 and pLM009, were integrated using X-ray irradiation. 100 L4 worms were irradiated with 5,000 rads using a MultiRad 350 (Precision X-Ray) then split onto 20 NGM plates. Plates were allowed to starve and then chunked to fresh plates, and allowed to starve again, repeating a total of three times. From each of the 20 plates, 10 worms that expressed the desired arrays were singled into 24-well plates for a total of 200 singled worms. The wells were screened three days later for homozygous integrants (100% of progeny expressed the co-transformation marker). This method was used to generate both *xnIs565 [lin-3p::YFP-PH, unc-119(+); lin-3p::exc-mScarlet, unc-119(+)]* and *xnIs569 [lin-3p::GFP1-10 + lin-3p::PH-mScarlet]*.

pLM008 was integrated separately using ‘FLInt’, as described (Malaiwong et al., 2023). pLM008 was co-injected (16 ng/µl) with *unc-119(+)* plasmid pJN254 (64ng/µl) (Nance et al., 2003) into strain EG7835 *[oxTi556 [eft-3p::tdTomato::H2B + Cbr-unc-119(+)]; unc-119(ed3)]* along with Cas9 protein (Berkeley), tracrRNA (IDT), and a crRNA (IDT) that cuts within tdTomato (Malaiwong et al., 2023). Non-red F1 were singled to individual plates, and plates with all non-red F2, ∼3/4 of which expressed pLM008 in L1 larvae, were saved. One insertion, *xnIs570*, was isolated.

pSA86 [*pgp-12p::mCherry*] was inserted in single copy at the *ttTi5605* Mos insertion site using MosSCI (Frokjaer-Jensen et al., 2008), yielding the *xnSi37* insertion.

### CRISPR/Cas9 genome editing

All edited alleles were created using CRISPR/Cas9 ribonucleoprotein complexes and repair DNA (ssDNA oligos or dsDNA PCR product) as described previously (Paix et al., 2015). Cas9 protein (Berkeley) was pre-incubated with crRNA (IDT) and tracrRNA (IDT), then mixed with repair DNA (ssDNA oligos or dsDNA PCR product) containing ∼35bp homology arms flanking the inserted sequence; repair templates contained silent mutations that prevented crRNA re-cutting of edited loci. The *dpy-10* co-CRISPR strategy was used to identify candidate edited worms. Dumpy and Roller F_1_ worms were singled and genotyped by PCR for the desired edit. crRNA sequences are listed in Table S2 and sequences of genes edited to insert fluorescent proteins or tags can be found in the Supplemental Methods.

### Immunostaining

Embryos were fixed and stained as described previously (Anderson et al. 2008). Primary antibodies used were: polyclonal rabbit anti-HMR-1 1:10,000 (Klompstra et al., 2015), monoclonal rat anti-GFP 1:1,000 (Nacalai Tesque), polyclonal rabbit anti-GFP 1:1,000 (Ab6556.25; AbCam), and monoclonal mouse anti-PSD-95 (Firestein and Rongo, 2001) (recognizes DLG-1) 1:200 (Affinity BioReagents). Secondary antibodies used were: Alexa Fluor 488 anti-rat IgG (Jackson ImmunoResearch), Alexa Fluor 647 anti-rabbit IgG (H+L) 1:200 (Invitrogen), Alexa Fluor 488 anti-rabbit IgG (H+L) 1:1,000 (Invitrogen), and Cy5 anti-mouse IgG (H+L) 1:200 (Jackson ImmunoResearch).

### Microscopy

For larval imaging experiments, L1 larvae were synchronized by cutting open ∼20 gravid adults in M9 buffer in a watch glass and allowing the released embryos to hatch overnight. L1 larvae were then immobilized using 5 µM levamisole and mounted on 4% agarose pads. Imaging was performed on a spinning disc confocal microscope (Nikon Eclipse Ti2, CSU-W1 or CSU-X1 spinning disk, Plan Fluor 40X 1.3 oil-immersion or Plan Apo 60x 1.2 NA water immersion lens, 488 nm and 561 nm lasers, and either Andor 888 Live EMCCD camera or matched Kinetix 22 sCMOS cameras (Teledyne) with a Twincam beam splitter. 0.3 µm spacing between Z stacks was used.

For embryonic imaging experiments, embryos were synchronized by cutting open ∼20-30 gravid adults in M9 in a watch glass. Comma-stage embryos were mounted via mouth pipette onto a 4% agar pad. Slides were placed in a humid chamber and the embryos were allowed to age for either 30 minutes or 1 hour depending on the developmental stage that was being examined. Imaging was then performed on a spinning disc confocal microscope (Nikon Eclipse Ti2, CSU-W1 spinning disk, 100X 1.35NA Silicone oil immersion objective, 488nm and 561nm lasers, matched Andor 888 Live EMCCD cameras). Timelapse images were acquired every 30 minutes.

### Image analysis

For excretory cell outgrowth measurements, ImageJ was used to measure the length of the posterior projections of the excretory cell in synchronized L1s. A line was drawn from the cell body of the excretory cell to the end of the posterior projections of the excretory cell in both WT and *ina-1* worms. Another line was drawn from the cell body of the excretory cell to the tip of the tail. The length of the excretory cell posterior projections was divided by the length of the embryo and multiplied by 100 to calculate the % outgrowth of the excretory cell.

To measure the epidermal basement membrane gap*, lam-2(qy20[lam-2::mNG]); xnSi37 [pgp12p::mCherry]* or *ina-1(xn219); lam-2(qy20[lam-2::mNG]); xnSi37 [pgp12p::mCherry]* L1 oriented with their dorsal or ventral sides up, and both left and right extensions of the excretory cell within five Z planes, were selected for image analysis. Using Fiji, an 8 µm line was drawn along the epidermal basement membrane, centered on the lateral excretory cell extension. The raw intensity of the basement membrane and excretory cell signal on the line was calculated after subtracting background fluorescence (determined for each marker by averaging an 8 µm line from a region outside of the worm). The LAM-2^mNG^ signal intensity was normalized by setting the average values of measurements in the first micron to a value of 1.0 A.U. (arbitrary units). mCherry signal intensity, which was used for orientation but not statistics, was normalized by setting the most intense value to 1.0 A.U. Microsoft Excel was used to generate graphs, to calculate the standard deviation of the LAM-2^mNG^ data, and to perform a two-tailed t-test comparing the last data point (8.0 µm) to all other data points except for those in the first micron used for normalization.

## Acknowledgements

We thank Matthew Buechner, Niels Ringstad, Meera Sundaram, and members of the Nance lab for insightful comments on the project or manuscript; Michael Cammer for invaluable input on imaging and image analysis; and Stephen Armenti and Mike Boxem for generously providing worm strains. Some strains were obtained from the *Caenorhabditis* Genetics Center (CGC), which is funded by NIH Office of Research Infrastructure Programs (P40 OD010440). Some experiments were performed in conjunction with the NYU Langone Microscopy Laboratory (RRID: SCR_017934) and the Genome Technology Center (RRID: SCR_017929). Funding for this study was provided by the National Institutes of Health through research grant R35GM118018 (J.N.) and training grant T32HD007520 (L.M.).

---

**Table S1.**
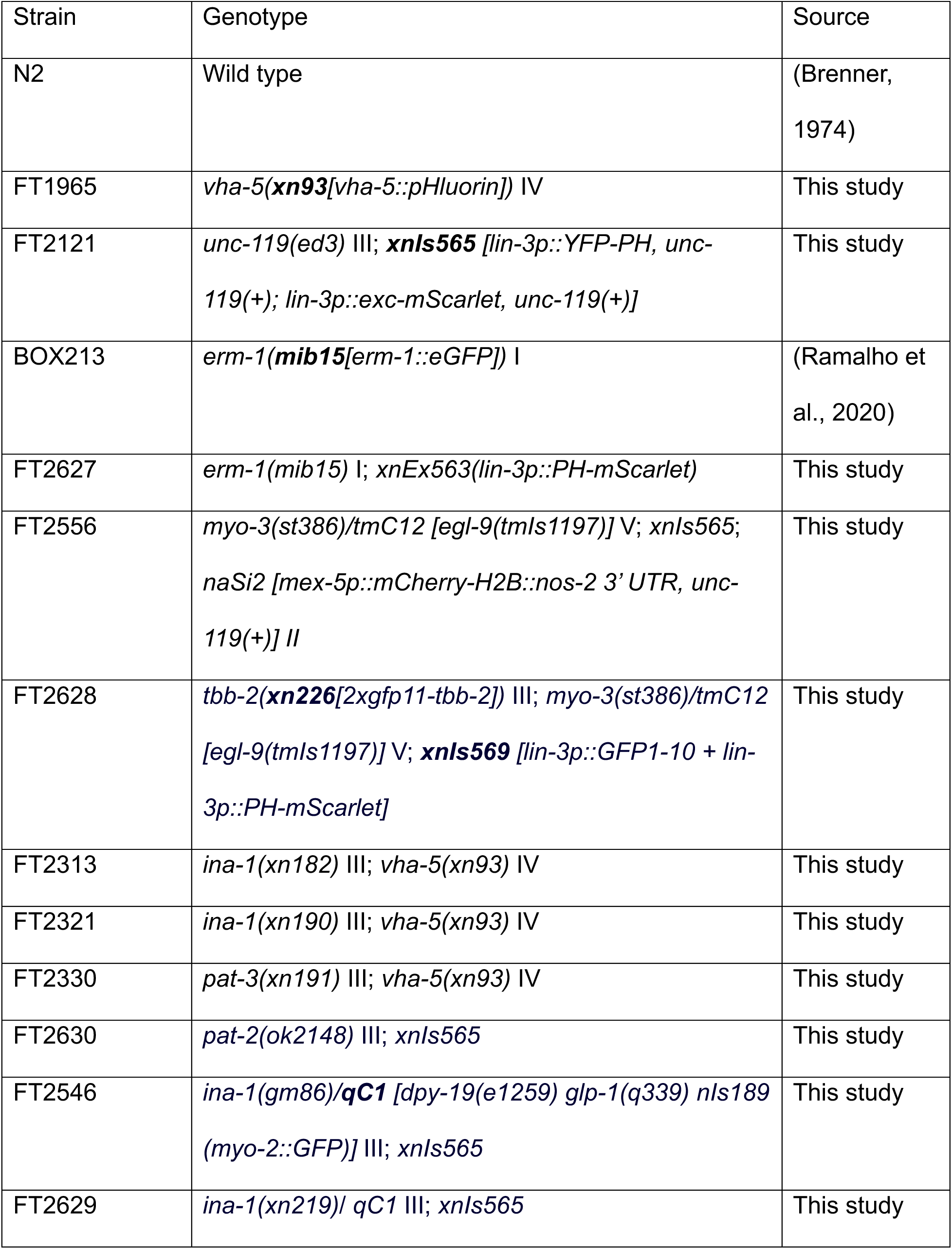
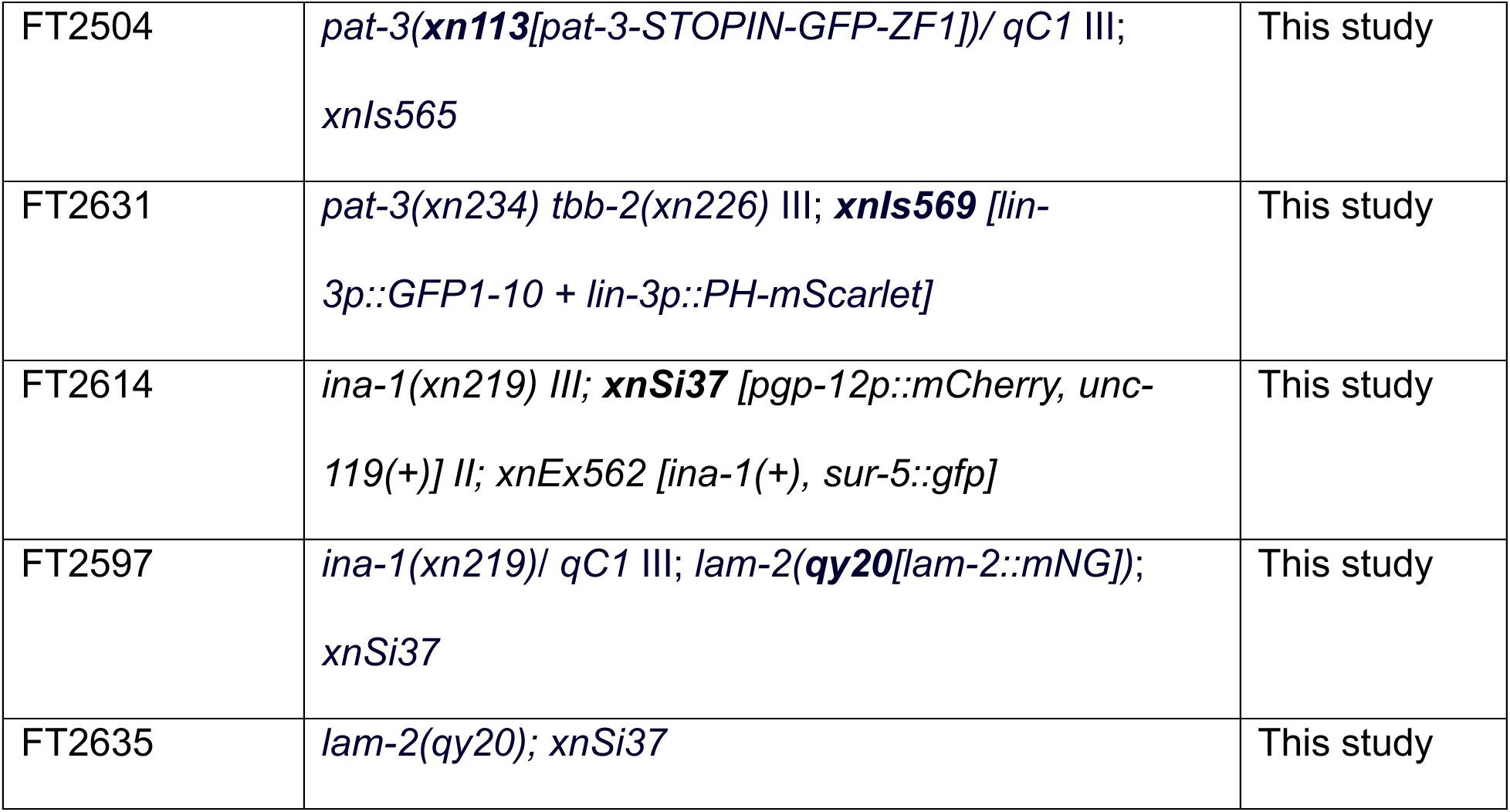
Worm strains.

**Table S2.**
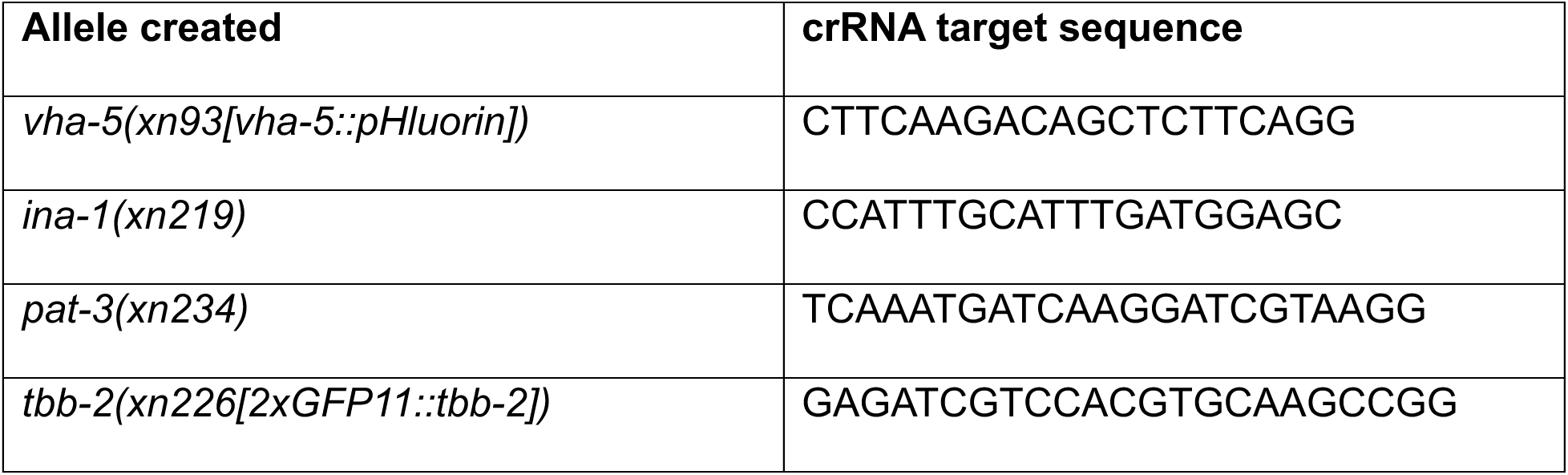
crRNAs used in CRISPR/Cas9 genome editing.

**Figure S1.**
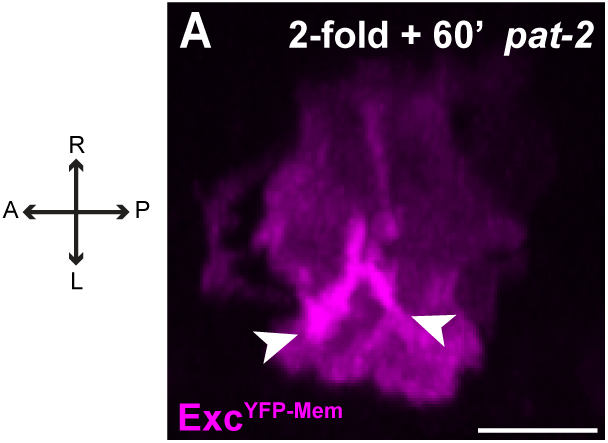
Excretory cell lumen branching in par-2 mutants. (A) Dorsal view of excretory cell and lumen growth and branching in a *par-2* mutant embryo expressing Exc^YFP-Mem^ at the indicated stage. Arrowheads indicate successful anterior-posterior branches. Compare to similarly staged *par-3* mutant embryo in Fig. 4D’ and *myo-3* embryo in Fig. 2A’’’. Scale bar, 5 µm.

## References

Abdus-Saboor, I., Mancuso, V. P., Murray, J. I., Palozola, K., Norris, C., Hall, D. H., Howell, K., Huang, K. and Sundaram, M. V. (2011). Notch and Ras promote sequential steps of excretory tube development in *C. elegans*. Development 138, 3545–3555.

Abrams, J. and Nance, J. (2021). A polarity pathway for exocyst-dependent intracellular tube extension. Elife 10, e65169.

Armenti, S. T., Chan, E. and Nance, J. (2014). Polarized exocyst-mediated vesicle fusion directs intracellular lumenogenesis within the *C. elegans* excretory cell. Dev Biol 394, 110–121.

Audhya, A., Hyndman, F., McLeod, I. X., Maddox, A. S., Yates, J. R., 3rd, Desai, A. and Oegema, K. (2005). A complex containing the Sm protein CAR-1 and the RNA helicase CGH-1 is required for embryonic cytokinesis in *Caenorhabditis elegans*. J Cell Biol 171, 267–279.

Banavar, S. P., Fowler, E. W. and Nelson, C. M. (2024). Biophysics of morphogenesis in the vertebrate lung. Curr Top Dev Biol 160, 65–86.

Baum, P. D. and Garriga, G. (1997). Neuronal migrations and axon fasciculation are disrupted in ina-1 integrin mutants. Neuron 19, 51–62.

Berry, K. L., Bulow, H. E., Hall, D. H. and Hobert, O. (2003). A *C. elegans* CLIC-like protein required for intracellular tube formation and maintenance. Science 302, 2134–2137.

Brenner, S. (1974). The genetics of Caenorhabditis elegans. Genetics 77, 71–94.

Buechner, M., Hall, D. H., Bhatt, H. and Hedgecock, E. M. (1999). Cystic canal mutants in *Caenorhabditis elegans* are defective in the apical membrane domain of the renal (excretory) cell. Dev Biol 214, 227–241.

Camelo, C. and Luschnig, S. (2021). Cells into tubes: Molecular and physical principles underlying lumen formation in tubular organs. Curr Top Dev Biol 143, 37–74.

Dong, B., Horie, T., Denker, E., Kusakabe, T., Tsuda, M., Smith, W. C. and Jiang, D. (2009). Tube formation by complex cellular processes in *Ciona intestinalis* notochord. Dev Biol 330, 237–249.

Firestein, B. L. and Rongo, C. (2001). DLG-1 is a MAGUK similar to SAP97 and is required for adherens junction formation. Mol Biol Cell 12, 3465–3475.

Frokjaer-Jensen, C., Davis, M. W., Hopkins, C. E., Newman, B. J., Thummel, J. M., Olesen, S. P., Grunnet, M. and Jorgensen, E. M. (2008). Single-copy insertion of transgenes in *Caenorhabditis elegans*. Nat Genet 40, 1375–1383.

Gavrilchenko, T., Simpkins, A. G., Simpson, T., Barrett, L. A., Hansen, P., Shvartsman, S. Y. and Schottenfeld-Roames, J. (2024). The *Drosophila* tracheal terminal cell as a model for branching morphogenesis. Proc Natl Acad Sci U S A 121, e2404462121.

Gervais, L. and Casanova, J. (2010). In vivo coupling of cell elongation and lumen formation in a single cell. Curr Biol 20, 359–366.

Gettner, S. N., Kenyon, C. and Reichardt, L. F. (1995). Characterization of beta pat-3 heterodimers, a family of essential integrin receptors in *C. elegans*. J Cell Biol 129, 1127–1141.

Gibson, D. G., Benders, G. A., Andrews-Pfannkoch, C., Denisova, E. A., Baden-Tillson, H., Zaveri, J., Stockwell, T. B., Brownley, A., Thomas, D. W., Algire, M. A., et al. (2008). Complete chemical synthesis, assembly, and cloning of a *Mycoplasma genitalium* genome. Science 319, 1215–1220.

Gobel, V., Barrett, P. L., Hall, D. H. and Fleming, J. T. (2004). Lumen morphogenesis in *C. elegans* requires the membrane-cytoskeleton linker *erm-1*. Dev Cell 6, 865–873.

Hagedorn, E. J., Yashiro, H., Ziel, J. W., Ihara, S., Wang, Z. and Sherwood, D. R. (2009). Integrin acts upstream of netrin signaling to regulate formation of the anchor cell’s invasive membrane in *C. elegans*. Dev Cell 17, 187–198.

Hedgecock, E. M., Culotti, J. G. and Hall, D. H. (1990). The *unc-5, unc-6*, and *unc-40* genes guide circumferential migrations of pioneer axons and mesodermal cells on the epidermis in *C. elegans*. Neuron 4, 61–85.

Hedgecock, E. M., Culotti, J. G., Hall, D. H. and Stern, B. D. (1987). Genetics of cell and axon migrations in *Caenorhabditis elegans*. Development 100, 365–382.

Herwig, L., Blum, Y., Krudewig, A., Ellertsdottir, E., Lenard, A., Belting, H. G. and Affolter, M. (2011). Distinct cellular mechanisms of blood vessel fusion in the zebrafish embryo. Curr Biol 21, 1942–1948.

Huang, C. C., Hall, D. H., Hedgecock, E. M., Kao, G., Karantza, V., Vogel, B. E., Hutter, H., Chisholm, A. D., Yurchenco, P. D. and Wadsworth, W. G. (2003). Laminin alpha subunits and their role in *C. elegans* development. Development 130, 3343–3358.

Hwang, B. J. and Sternberg, P. W. (2004). A cell-specific enhancer that specifies lin-3 expression in the *C. elegans* anchor cell for vulval development. Development 131, 143–151.

Hynes, R. O. and Zhao, Q. (2000). The evolution of cell adhesion. J Cell Biol 150, F89–96.

Joseph, B. B., Blouin, N. A. and Fay, D. S. (2018). Use of a Sibling Subtraction Method for Identifying Causal Mutations in *Caenorhabditis elegans* by Whole-Genome Sequencing. G3 (Bethesda) 8, 669–678.

Kanchanawong, P. and Calderwood, D. A. (2023). Organization, dynamics and mechanoregulation of integrin-mediated cell-ECM adhesions. Nature reviews. Molecular cell biology 24, 142–161.

Kao, G., Huang, C. C., Hedgecock, E. M., Hall, D. H. and Wadsworth, W. G. (2006). The role of the laminin beta subunit in laminin heterotrimer assembly and basement membrane function and development in *C. elegans*. Dev Biol 290, 211–219.

Keeley, D. P., Hastie, E., Jayadev, R., Kelley, L. C., Chi, Q., Payne, S. G., Jeger, J. L., Hoffman, B. D. and Sherwood, D. R. (2020). Comprehensive Endogenous Tagging of Basement Membrane Components Reveals Dynamic Movement within the Matrix Scaffolding. Dev Cell 54, 60–74 e67.

Klompstra, D., Anderson, D. C., Yeh, J. Y., Zilberman, Y. and Nance, J. (2015). An instructive role for *C. elegans* E-cadherin in translating cell contact cues into cortical polarity. Nat Cell Biol 17, 726–735.

Kolotuev, I., Hyenne, V., Schwab, Y., Rodriguez, D. and Labouesse, M. (2013). A pathway for unicellular tube extension depending on the lymphatic vessel determinant Prox1 and on osmoregulation. Nat Cell Biol 15, 157–168.

Lenard, A., Ellertsdottir, E., Herwig, L., Krudewig, A., Sauteur, L., Belting, H. G. and Affolter, M. (2013). In vivo analysis reveals a highly stereotypic morphogenetic pathway of vascular anastomosis. Dev Cell 25, 492–506.

Levi, B. P., Ghabrial, A. S. and Krasnow, M. A. (2006). *Drosophila* talin and integrin genes are required for maintenance of tracheal terminal branches and luminal organization. Development 133, 2383–2393.

Liegeois, S., Benedetto, A., Garnier, J. M., Schwab, Y. and Labouesse, M. (2006). The V0-ATPase mediates apical secretion of exosomes containing Hedgehog-related proteins in *Caenorhabditis elegans*. J Cell Biol 173, 949–961.

Malaiwong, N., Porta-de-la-Riva, M. and Krieg, M. (2023). FLInt: single shot safe harbor transgene integration via Fluorescent Landmark Interference. G3 (Bethesda) 13.

McIntyre, D. C. and Nance, J. (2023). Niche cells regulate primordial germ cell quiescence in response to basement membrane signaling. Development 150.

Mello, C. C., Kramer, J. M., Stinchcomb, D. and Ambros, V. (1991). Efficient gene transfer in *C. elegans*: extrachromosomal maintenance and integration of transforming sequences. The EMBO journal 10, 3959–3970.

Nance, J., Munro, E. M. and Priess, J. R. (2003). *C. elegans* PAR-3 and PAR-6 are required for apicobasal asymmetries associated with cell adhesion and gastrulation. Development 130, 5339–5350.

Nelson, F. K., Albert, P. S. and Riddle, D. L. (1983). Fine structure of the *Caenorhabditis elegans* secretory-excretory system. Journal of ultrastructure research 82, 156–171.

Nelson, F. K. and Riddle, D. L. (1984). Functional study of the *Caenorhabditis elegans* secretory-excretory system using laser microsurgery. J Exp Zool 231, 45–56.

Noma, K., Goncharov, A., Ellisman, M. H. and Jin, Y. (2017). Microtubule-dependent ribosome localization in *C. elegans* neurons. Elife 6.

Packer, J. S., Zhu, Q., Huynh, C., Sivaramakrishnan, P., Preston, E., Dueck, H., Stefanik, D., Tan, K., Trapnell, C., Kim, J., et al. (2019). A lineage-resolved molecular atlas of *C. elegans* embryogenesis at single-cell resolution. Science 365.

Paix, A., Folkmann, A., Rasoloson, D. and Seydoux, G. (2015). High Efficiency, Homology-Directed Genome Editing in Caenorhabditis elegans Using CRISPR-Cas9 Ribonucleoprotein Complexes. Genetics 201, 47–54.

Ramalho, J. J., Sepers, J. J., Nicolle, O., Schmidt, R., Cravo, J., Michaux, G. and Boxem, M. (2020). C-terminal phosphorylation modulates ERM-1 localization and dynamics to control cortical actin organization and support lumen formation during *Caenorhabditis elegans* development. Development 147.

Rasmussen, J. P., English, K., Tenlen, J. R. and Priess, J. R. (2008). Notch signaling and morphogenesis of single-cell tubes in the *C. elegans* digestive tract. Dev Cell 14, 559–569.

Shaye, D. D. and Greenwald, I. (2015). The disease-associated formin INF2/EXC-6 organizes lumen and cell outgrowth during tubulogenesis by regulating F-actin and microtubule cytoskeletons. Dev Cell 32, 743–755.

Shaye, D. D. and Soto, M. C. (2021). Epithelial morphogenesis, tubulogenesis and forces in organogenesis. Curr Top Dev Biol 144, 161–214.

Smith, H. E., Fabritius, A. S., Jaramillo-Lambert, A. and Golden, A. (2016). Mapping Challenging Mutations by Whole-Genome Sequencing. G3 (Bethesda) 6, 1297–1304.

Soulavie, F., Hall, D. H. and Sundaram, M. V. (2018). The AFF-1 exoplasmic fusogen is required for endocytic scission and seamless tube elongation. Nat Commun 9, 1741.

Stone, C. E., Hall, D. H. and Sundaram, M. V. (2009). Lipocalin signaling controls unicellular tube development in the *Caenorhabditis elegans* excretory system. Dev Biol 329, 201–211.

Sulston, J. E., Schierenberg, E., White, J. G. and Thomson, J. N. (1983). The embryonic cell lineage of the nematode *Caenorhabditis elegans*. Dev Biol 100, 64–119.

Sundaram, M. V. and Buechner, M. (2016). The *Caenorhabditis elegans* Excretory System: A Model for Tubulogenesis, Cell Fate Specification, and Plasticity. Genetics 203, 35–63.

Sundaram, M. V. and Cohen, J. D. (2017). Time to make the doughnuts: Building and shaping seamless tubes. Seminars in cell & developmental biology 67, 123–131.

Waterston, R. H. (1989). The minor myosin heavy chain, mhcA, of *Caenorhabditis elegans* is necessary for the initiation of thick filament assembly. The EMBO journal 8, 3429–3436.

White, J. G., Southgate, E., Thomson, J. N. and Brenner, S. (1986). The structure of the nervous system of the nematode *Caenorhabditis elegans*. Philos Trans R Soc Lond B Biol Sci 314, 1–340.

Williams, B. D. and Waterston, R. H. (1994). Genes critical for muscle development and function in *Caenorhabditis elegans* identified through lethal mutations. J Cell Biol 124, 475–490.

Yochem, J. (2006). Nomarski images for learning the anatomy, with tips for mosaic analysis. In WormBook : the online review of C. elegans biology (ed. D. Fay): The C. elegans Research Community.

Yochem, J., Gu, T. and Han, M. (1998). A new marker for mosaic analysis in *Caenorhabditis elegans* indicates a fusion between hyp6 and hyp7, two major components of the hypodermis. Genetics 149, 1323–1334.

Yurchenco, P. D. and Kulczyk, A. W. (2024). Polymerizing laminins in development, health, and disease. J Biol Chem 300, 107429.

Zhao, Z., Fang, L., Chen, N., Johnsen, R. C., Stein, L. and Baillie, D. L. (2005). Distinct regulatory elements mediate similar expression patterns in the excretory cell of *Caenorhabditis elegans*. J Biol Chem 280, 38787–38794.

